# Genetic overlap between in-scanner head motion and the default network connectivity

**DOI:** 10.1101/087023

**Authors:** Yuan Zhou, Jie Chen, Yu L.L. Luo, Dang Zheng, Li-Lin Rao, Xinying Li, Jianxin Zhang, Shu Li, Karl Friston, Xi-Nian Zuo

## Abstract

The association between in-scanner head motion and intrinsic functional connectivity (iFC) may confound explanations for individual differences in functional connectomics. However, the etiology of the correlation between head motion and iFC has not been established. This study aimed to investigate genetic and environmental contributions on the association between head motion and iFC using a twin dataset (175 same-sex twin pairs, aged 14-23 years, 48% females). After establishing that both head motion and default network iFC are moderately heritable, we found large genetic correlations (-0.52 to -0.73) between head motion and the default network iFCs. Common genes can explain 48% - 61% of the negative phenotypic correlation between the two phenotypes. These results advance our understanding of the relationship between head motion and iFC, and may have profound implications for interpreting individual differences in default network connectivity in clinical research and brain-behavior association.

## Introduction

Individual differences in how people think and behave are rooted in the variability of the brain’s functional anatomy (Mueller, et al., 2013). Intrinsic functional connectivity (iFC) – derived from resting-state functional magnetic resonance imaging (rfMRI) – offers a powerful approach to map functional brain architectures (Fox and Raichle, 2007). Individual differences in iFC have been demonstrated to be stable across time (Zuo and Xing, 2014), and can usefully characterize inter-individual variability in healthy participants as well as provide diagnostic correlates of symptom severity in neuropsychiatric disorders (Broyd, et al., 2009; Finn, et al., 2015; Jiang, et al., 2013; Kelly, et al., 2012; Tavor, et al., 2016; Zhang and Raichle, 2010). Recently, one particular concern about the phenotypic correlate of iFC has focused on in-scanner head motion: three groups have reported that head motion can obscure differences in short-range iFC, relative to long-range correlations (Power, et al., 2012; Satterthwaite, et al., 2012; Van Dijk, et al., 2012). This motion-iFC relationship seems to challenge the interpretation of associations of iFC with age (Power, et al., 2012; Satterthwaite, et al., 2012) and neuropsychiatric disorders (e.g., autism) (Deen and Pelphrey, 2012), because younger (and e.g., autistic) subjects tend to move more in the scanner (Deen and Pelphrey, 2012; Power, et al., 2015). Therefore, it is crucial to understand the mechanisms underlying the motion-iFC relationship.

The prevalent view is that the motion-iFC association is an artifact of in-scanner head motion (Power, et al., 2012; Power, et al., 2015; Satterthwaite, et al., 2012; Van Dijk, et al., 2012). However, other explanations have been proposed recently. Firstly, the motion-iFC associations – located specifically in sensorimotor and visual cortex (positive correlations) as well as the default network (negative correlations) – were found in highly compliant young adults, suggesting that part of the motion-iFC association might be mediated neuronally by functional segregated brain systems (Pujol, et al., 2014). Furthermore, after minimizing motion-related artifacts, reduced long-range iFC primarily within the default network was found to correlate with head motion in inter-individual analysis, but not in analysis of intra-subject data. This suggests that the iFC in the default network might represent a neurobiological trait that predisposes some individuals to head movements (Zeng, et al., 2014). Moreover, the repeatedly observed correlation between head motion and iFC of the default network (Pujol, et al., 2014; Satterthwaite, et al., 2012; Van Dijk, et al., 2012; Zeng, et al., 2014) is stable within individuals across time (Zeng, et al., 2014) and shows high test-retest reliability (Yan, et al., 2013a). Collectively, these observations suggest the association between head motion and iFC, particularly that between small movements and iFC of the default network, may reflect a trait-like property that cannot be fully explained as an artifact of head motion. This suggests that there are alternative mechanisms underlying the association.

Here, we set out to understand the motion-iFC association from the perspective of genetic etiology. We hypothesized that intersubject variability in iFC, particular within the default network, and the tendency to move during scanning could be partly driven by common genetic factors. This genetic overlap hypothesis is proposed based on several recent observations on the heritability of both head motion and iFCs. On one hand, a moderate heritability of the in-scanner head motion has been reported (Couvy-Duchesne, et al., 2014; Couvy-Duchesne, et al., 2016). On the other hand, the iFC in the default network has shown moderate test-retest reliability (Yan, et al., 2013a; Zuo and Xing, 2014) and heritability (Fu, et al., 2015; Glahn, et al., 2010; Korgaonkar, et al., 2014; Yang, et al., 2016). To test our hypothesis, we employed a twin design (Neale and Cardon, 1992) to determine the genetic and environmental influences on the motion-iFC correlation. Specifically, we used bivariate genetic models to estimate the extent of genetic (*r_g_*) and environmental correlation (*r_e_*) between head motion and iFC (Fig. S1 in supplementary materials). We were particularly interested in the default network, because the cognitive functions subserved by the default network (such as perspective taking, introspection and mind-wandering) (Mason, et al., 2007) may provide a potential account of the genetic link between the iFC of default network and the tendency to head motion. In other words, subjects with more evident default network functional connectivity may engage more self-referential (less reactive) neuronal processing and consequently move less.

## Materials and Methods

### Participants

The sample was based on 215 same-sex twin pairs (116 monozygotic and 99 dizygotic pairs) from the Beijing Twin Study (BeTwiSt) at the Institute of Psychology, Chinese Academy of Sciences **(Chen, et al., 2013)**. Age range of the sample was 14-23 years (mean = 17.4 years; 49.3% female). The BeTwiSt twins are largely representative of the general youth population in Beijing **(Chen, et al., 2013)**. The twin zygosity was determined by DNA analyses for 210 pairs and a questionnaire method for five pairs **(Chen, et al., 2010)**, who were excluded for the data analyses. After exlcuding 35 twins pairs showing less “good” volumes data (number = 33) or showing outlier in the headmotion measure (> 3 interquartile ranges from the sample median) (number = 2), 175 same-sex twin pairs ((100 monozygotic and 75 dizygotic pairs; age = 17.4 ± 2.0 years; 48% females) were recruited into the final data analyses.

No participants had any history of psychiatric diagnoses, or any neurological or metabolic illnesses. Written informed consent has been obtained from all participants or their guardians. This study was approved by the Institutional Review Board of the Institute of Psychology of the Chinese Academy of Sciences and the Institutional Review Board of Beijing MRI Center for Brain Research.

### Data Acquisition

The MRI data were acquired with a 3.0-Tesla Siemens MRI scanner (MAGNETOM TRIO) in the Bejing MRI Center for Brain Research. Whole-brain rfMRI scans were collected in 32 axial slices using an echo-planar imaging (EPI) sequence (repetition time [TR] = 2000 ms, echo time [TE] = 30 ms; flip angle [FA] = 90°, matrix = 64 × 64; field of view [FoV] = 220 × 220 mm^2^; slice thickness = 3 mm; slice gap = 1 mm; 180 volumes). During the rfMRI acquisition, the participants were explicitly instructed to lie supine, stay relaxed with their eyes closed, and move as little as possible. High-resolution T1-weighted images were acquired in a sagittal orientation using a magnetization-prepared rapid gradient-echo (MPRAGE) sequence (TR/TE = 2530/3.37 ms; FA = 7^o^; FoV = 240 mm^2^; 1-mm in-plane resolution; slice thickness = 1.33 mm, no gap; 144 slices).

### Functional imaging preprocessing

Conventional functional imaging preprocessing was performed using the Data Processing Assistant for Resting-State fMRI (DPARSF 4.1, http://www.restfmri.net) (Yan, et al., 2016), including the removal of the first 10 volumes, corrections for slice timing and spatial registration, nuisance variable regression, spatial normalization with 3-mm cubic voxels, spatial smooth of 6 mm FWHM and temporal band-pass (0.01–0.1 Hz) filtering. The nuisance variables include 24 motion parameters (6 head motion parameters, 6 head motion parameters one time point before, and the 12 corresponding squared items), the signal averaged over the individual segmented CSF and white matter (WM) regions, linear and quadratic trends (Yan, et al., 2013b).

### Head motion measurements and scrubbing

The volume-based frame-wise displacement (FD) was used to quantify head motion (Power, et al., 2012; Satterthwaite, et al., 2012; Van Dijk, et al., 2012). To further reduce the influences of motion-related artifacts on rfMRI-derived iFC, we employed the volume-based scrubbing regression by including scrubbing regressors into the multiple linear regression (Yan, et al., 2013a). The time points with a threshold of FD > 0.2 mm as well as 1 back and 2 forward frames were identified and then modeled as a separate regressor in the regression model of realigned rfMRI data. We set a 3-minute criterion and excluded subjects who had less than 90 “good” volumes data (Yan, et al., 2013a).

### Distant functional connectivity measured by long-range degree centrality

The network degree centrality (DC) (Sepulcre, et al., 2010) was employed to examine iFC. For each twin, the correlations between the time courses of each pair of gray matter voxels were calculated and used to construct an undirected adjacency matrix by thresholding ensuing correlations at r > 0.25 (Buckner, et al., 2009; Zhou, et al., 2014). An arbitrary distance threshold (16 mm) was used to identify long connections; the threshold was determined according the smoothing kernel applied to our data and known anatomical connectivity (Gilbert and Wiesel, 1989). The degree of distant iFC was computed as the sum of the weights of the connections for each voxel outside its immediate neighborhood (defined as a 16 mm radius sphere). The individual voxel-wise degree maps were converted into Z-score maps by subtracting the mean iFC across the entire brain and dividing by the standard deviation of the whole-brain iFC (Yan, et al., 2013b; Zuo, et al., 2012). This distance-dependent centrality computation has been included in a new release of the Connectome Computation System (CCS: https://github.com/zuoxinian/CCS) (Xu, et al., 2015).

Multiple regression analysis was performed to examine the correlations between the iFC and head motion (quantified using Power’s mean FD) while accounting for the effects of age and gender. Statistical significance was set at P < 0.05 using FWE to correct for multiple comparisons.

### Univariate genetic analyses

We firstly compared ICC for MZ and DZ twins. When MZ twins correlate more highly than DZ twins, genetic influences (A) are implied. We then employed the univariate genetic modeling to estimate the ACE parameters (full model) and their confidence intervals using the OpenMx package (http://openmx.psyc.virginia.edu). For the head motion measure and voxel-wise iFC, we first regressed out the age and gender effects and then submitted the residuals to univariate modelling. Path coefficients were estimated using the maximum likelihood method and the goodness of model fit was indicated by the -2 times the log likelihood (-2LL). The relative fit of nested models was tested by removing paths from the full model and assessing the significance of removal using a χ^2^ test. A significant χ^2^ difference resulted in rejecting the submodel, otherwise the more parsimonious submodels were accepted (Bollen, 1989). We also used the Akaike information criterion (AIC) (Akaike, 1974) and the Bayesian information criterion (BIC) (Schwarz, 1978), where lower values indicating greater model evidence. We used the posterior probability maps (PPMs) to identify the regions showing genetic effect at ≥90% confidence level (Friston and Penny, 2003).

### Bivariate genetic analyses

Bivariate genetic analysis – an extension of the univariate analysis above – was used to partition the covariance between two phenotypes into A, C and E components. The central parameter in bivariate genetic model is the genetic correlation (*r*_g_), which indicates the extent to which the two phenotypes are genetically mediated. We further calculated the proportion of phenotypic correlation (rph) due to correlated genetic effects A (rph-a = a_1_×*r*_*g*_×a_2_), and correlated non-shared environmental effects E (rph-e = e_1_×*r*_e_×*e*_2_) expressed as proportions of rph (Fig. S1.A). An alternative representation of bivariate genetic modeling is Cholesky decomposition (Fig. S1.B), which calculates how much of the heritability or environmental effect of one phenotype can be attributed to another. For bivariate modelling, we focused on those clusters showing significant heritability at ≥90% confidence level (and having at least 5 voxels). The clusters were extracted and the averaged degree of distant iFC was subject to bivariate genetic analysis.

### Trivariate genetic modeling of global signal, head motion and brain connectivity

The global mean signal was extracted during preprocessing. The global signal (GS) was estimated by averaging rfMRI time series across all voxels within the whole brain and its amplitude of low-frequency fluctuations (GSalff) was calculated. A univariate genetic modeling analysis was firstly performed to test whether the global signal is heritable. Then trivariate genetic modeling analyses were performed to explore the genetic correlations among the GS, head motion and functional connectivity.

## Results

The novel contribution of the present work is to establish a significant genetic correlation between the default network iFC and in-scanner head motion. Specifically, common genes accounted for more than half of the negative phenotypic correlation between head motion and default network iFC. Multivariate genetic analyses showed that the genetic correlations between head motion and default network connectivity remained significant, controlling for their genetic overlap with global (whole brain) signals.

### Phenotypic correlation between head motion and functional connectivity

In our sample of 175 same-sex twin pairs, the average head motion measured by the volume-based frame-wise displacement (FD) (Power, et al., 2012) was 0.13 mm with a standard deviation of 0.05. We then assessed the phenotypic correlation between head motion and iFC characterized by voxel-wise distant degree centrality. The degree of distant connectivity was calculated for each voxel by counting the number of voxels outside the immediate neighborhood (a sphere with the radius of 16 mm) (details see Methods) (Gilbert and Wiesel, 1989).

Significant motion-iFC correlations (family-wise error, FWE corrected *P* < 0.05) were dominated by negative ones within the default network (Fig. 1), while positive motion-iFC correlations were largely restricted to subcortical regions and voxels close to white matter (Table S1 in supplementary materials). Specifically, moderate negative motion-iFC correlations (Fig. 1A) were detected in the bilateral posterior cingulate cortex (PCC), medial prefrontal cortex (MPFC), angular gyri (AG), and anterior temporal cortex, which are the default network components according to an unbiased whole-brain network parcellation (Fig. 1B) (Yeo, et al., 2011). Moderate positive motion-iFC correlations were observed in the bilateral caudate, the right temporal cortex, and the right cerebellum (FWE, corrected *P* < 0.05).

**Fig. 1.**
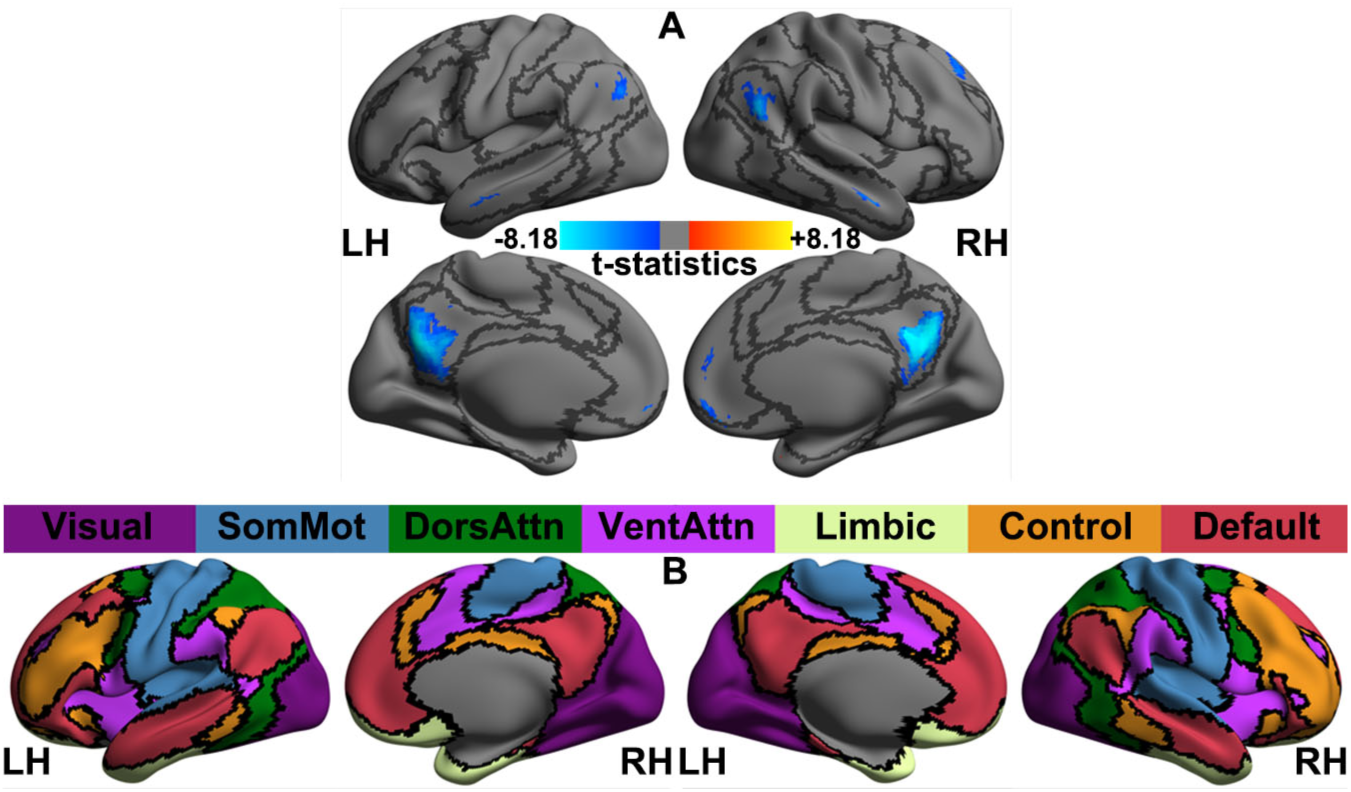
Phenotypic correlation between in-scanner head motion and intrinsic functional connectivity (iFC) measured by the degree of distant connectivity (A). A large-scale cortical network parcellation derived from 1000 healthy participants’ rfMRI images in (24) is shown in (B) as a reference for the locations of seven common brain networks, including visual (Visual), somatomotor (SomMot), dorsal attention (DorsAttn), ventral attention (VentAttn), limbic (Limbic), frontoparietal control (Control) and default (Default) networks. Statistical maps are rendered onto the template surface ‘32k_ConteAtlas’ provided in the Workbench Connectome pipeline from the Human Connectome Project by using the CCS visualization module, with lateral and medial views for both the left (LH) and right hemispheres (RH). Black curves indicate the network boundaries.

### Univariate genetic analyses

For the in-scanner head motion measure, the intraclass correlation coefficient (ICC), applied to the cross-twin correlation, was significantly higher in monozygotic twins (MZ) (0.45) than dizygotic twins (DZ) (-0.05) (Fisher’s Z test, one-tailed, *P* < 0.001), suggesting genetic influences. We then quantified the additive genetic (A), shared- (C), and non-shared environmental (E) contributions to the variance of head motion with the univariate genetic model (Plomin, et al., 2013). As the shared environmental effect (C) equaled 0, the AE model was fitted (Table S2), resulting in heritability of 25% (95% CI: 6%-42%) and non-shared environmental effect of 75% (95% CI: 58%-94%).

For the iFC negatively correlated with motion, the ICC_MZ_ were generally higher than the ICC_DZ_ (MZ: median = 0.32, interquartile range = 0.18; DZ: median = 0.20, interquartile range = 0.21) (Fig. 2A and 2B). The univariate genetic analysis was then performed for voxels showing genetic influences (see above). We compared the AE and E models because the C estimates were very close to or equal 0 in the preliminary analyses (results available on request). We derived posterior probability maps (PPMs) (Friston and Penny, 2003) to identify brain regions with at least 90% confidence that the AE model was better than the E model. We found that the distant iFCs in the bilateral PCC, the MPFC and the left AG were significantly heritable (Fig. 3 and Table S3). Independent univariate genetic analyses showed that the averaged iFC in these regions were moderately heritable (31%-39%) (Table S4). However, for regions showing positive motion-iFC correlations, no significant heritability was found. A whole brain analysis of the variance estimates for the distant iFC further endorsed the finding that default network iFCs are significantly heritable (Fig. S2).

**Fig. 2.**
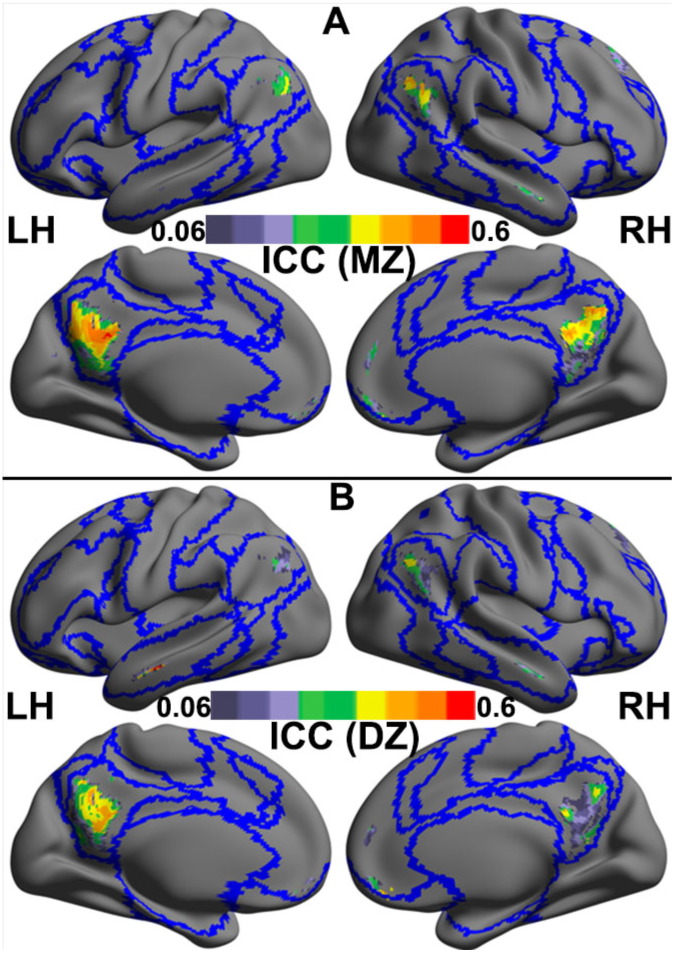
Cross-twin correlations of intrinsic functional connectivity in the regions whose degree of distant functional connectivity negatively correlated with in-scanner motion head motion are shown for monozygotic (MZ) twins (A) and dizygotic (DZ) twins (B). These correlations are estimated as the intra-class correlation coefficient (ICC) are rendered onto the template surface ‘32k_ConteAtlas’ provided in the Workbench Connectome pipeline from the Human Connectome Project using CCS, with lateral and medial views for both the left (LH) and right hemispheres (RH). Blue curves indicate the boundaries among the Yeo’s seven common networks as depicted in Fig. 1B.

**Fig. 3.**
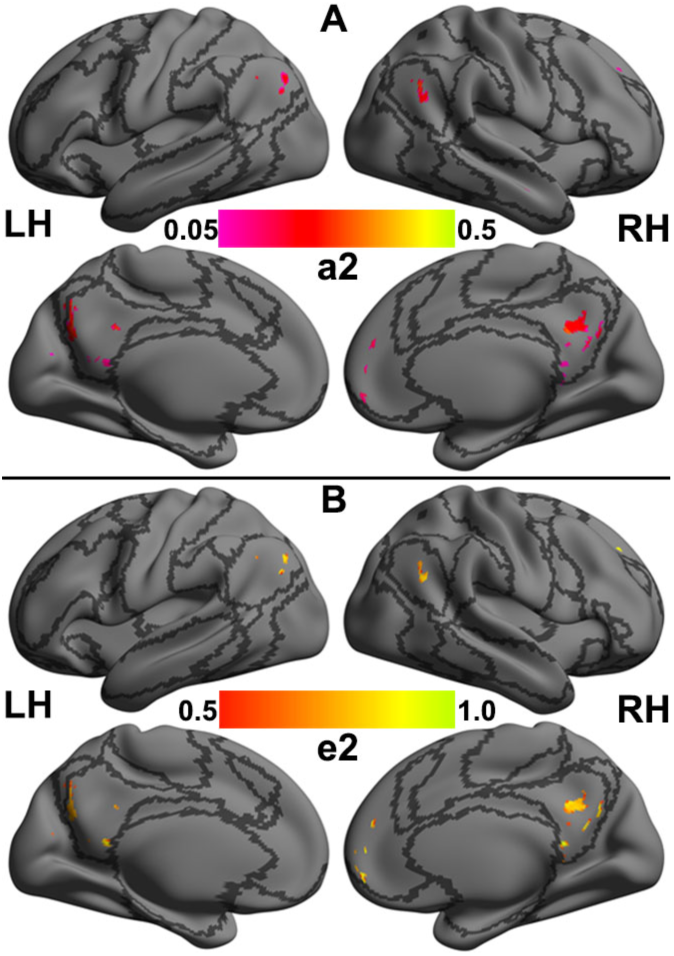
Variance components estimated for intrinsic functional connectivity (iFC) measured by the degree of distant functional connectivity of the regions showing significant negative correlations between their iFC and head motion: A) genetic variance (a^2^), B) non-shared environmental variance (e^2^). These variance maps are rendered onto the template surface ‘32k_ConteAtlas’ provided in the Workbench Connectome pipeline from the Human Connectome Project using the CCS visualization module, with lateral and medial views for both the left (LH) and right hemispheres (RH). Black curves indicate the boundaries among the Yeo’s seven common networks as depicted in Fig. 1B.

### Bivariate genetic analyses

Based on the results of univariate genetic analyses, our bivariate analyses focused on head motion and its negatively correlated iFC (i.e., distant iFC of default network). The cross-twin cross-trait correlations suggest genetic influences on the motion-iFC correlation (Table S5). We then used the bivariate genetic analysis to partition the covariance between the head motion and iFC into A, C and E components (Fig. S1.A). The genetic correlation (*r_g_*) between head motion and the iFC of the bilateral PCC, MPFC and left AG were significant and large, ranging from -0.52 to -0.73 (Fig. 4A, and Table S6). The proportions of the phenotypic association accounted for by overlapping genetic factors (rph-a) were moderate to large (48% to 61%) (Fig. 4B, and Table S6).

**Fig. 4.**
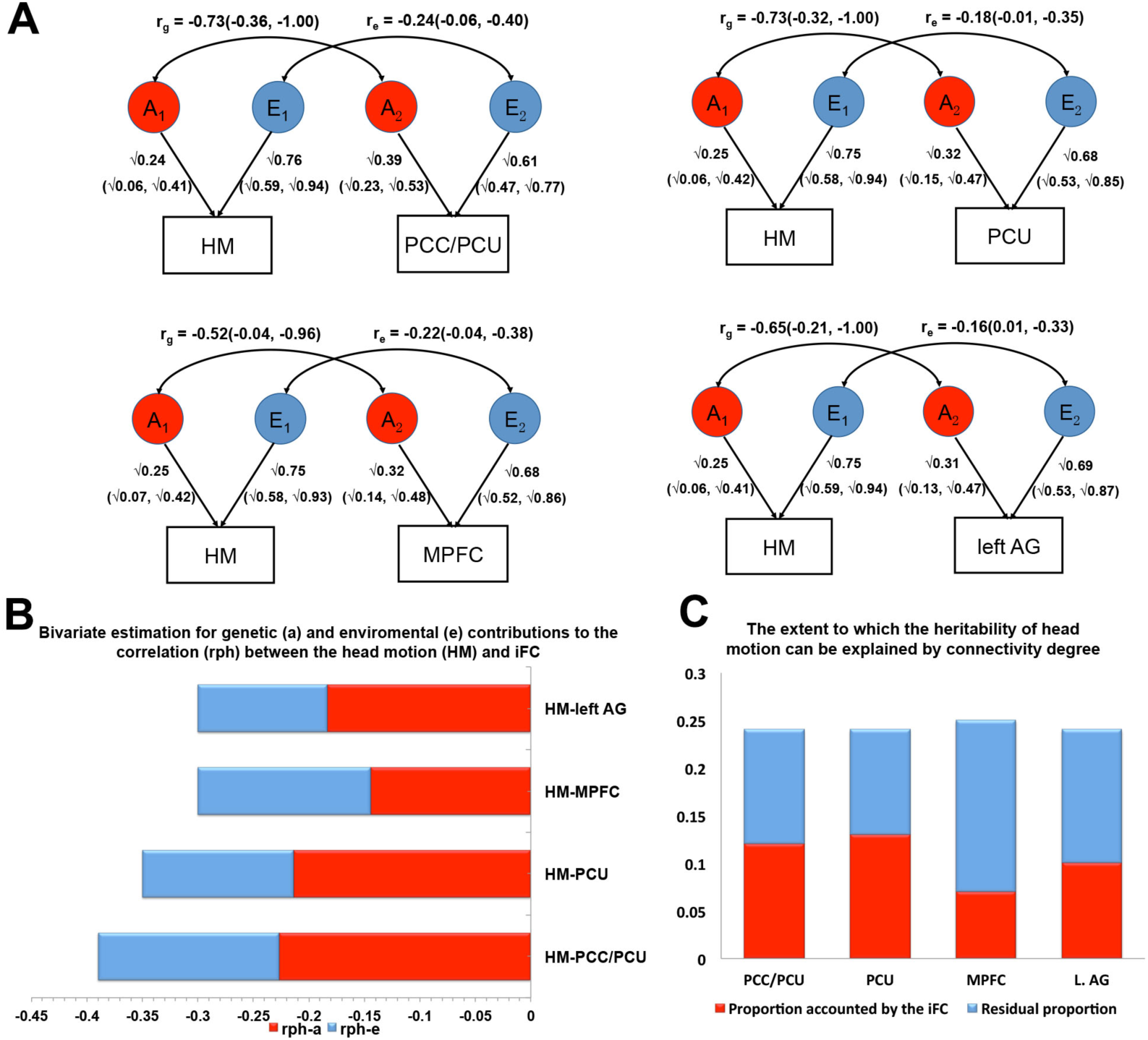
Bivariate genetic/environmental correlation modeling of head motion and intrinsic functional connectivity (iFC) measured by the degree of distant functional connectivity. The genetic (*r*_g_) and environmental (*r_e_*) correlation between the head motion and the iFC were shown in (A). The bivariate estimation for genetic (rph-a) and environmental (rph-e) contributions to the correlation between the head motion and iFC (the length of the bar) were shown in (B). The extent to which the heritability of the head motion (the height of the bar) can be explained by the iFC was shown in (C).

We further conducted the Cholesky analyses (Fig. S1.B) to separate the heritability of head motion into variance attributed to (or shared with) the iFC and residual variance. The results showed that the proportion of heritability on head motion explained by iFC was moderate to large (28% to 54%) (Fig. 4C and Table S7); the additive accounting of the four iFC of default network can reach 58%.

### Trivariate genetic modeling of global signal, head motion and brain connectivity

To exclude the possible confounding of global signal on the genetic association between head motion and distant iFC of default network, we conducted the trivariate genetic modeling of head motion, distant iFC of default network and global signal. We used the mean amplitude of low-frequency (ALFF) of rfMRI time series across all voxels of whole brain as the indicator of global signal. We found that the heritability of global signal was 39% (Table S8). Trivariate genetic analysis showed that the magnitudes genetic correlations between the distant iFC of the default network and the head motion are unchanged (i.e., same to the bivariate analyses), after controlling their genetic overlaps with global signal (Table S9). These results suggest that the genetic overlap between head motion and distant iFC of default network is not confounded by global signal.

## Discussion

We found a significant genetic correlation between the tendency to in-scanner head motion and default network functional connectivity. This finding advances our understanding the relationship between head motion and intrinsic functional connectivity and augments the traditional notion that head motion causes functional connectivity artifact. These conclusions may have substantial implications for the interpretation of rfMRI studies.

We found that in-scanner head motion was positively correlated with the bilateral caudate, the right temporal cortex and cerebellum, and primarily negatively correlated with the iFC of core default network regions (Andrews-Hanna, et al., 2010b), such as the bilateral PCC and MPFC. These negative phenotypic correlations are compatible with previous findings (Pujol, et al., 2014; Satterthwaite, et al., 2012; Van Dijk, et al., 2012; Zeng, et al., 2014), suggesting that the negative motion-iFC association within default network is robust. Univariate genetic analyses revealed that the tendency to head motion is moderately heritable (25%), which is consistent with the previously reported moderate heritability of head motion measures (16, 19). These findings indicate that, rather than an artifact, in-scanner head motion partly reflects a biological trait. For motion-related iFC measures, we found significant genetic influences mainly in the default network regions, including the bilateral PCC, MPFC, the left angular gyrus. This is in good agreement with previous findings regarding the genetic basis of default network connectivity (Glahn, et al., 2010; Korgaonkar, et al., 2014; Yang, et al., 2016). However, we did not find genetic influences in regions whose functional connectivity positively correlated with head motion.

Beyond these replications, a novel finding in this study is that the motion-iFC correlations are largely accounted for by common genetic etiology. Specifically, the genetic correlations between head motion and default network connectivity are significant and large (-0.52 to -0.73), suggesting that a large part of genes influencing default network connectivity also underlie the tendency of head motion. More than half of the heritability of head motion (i.e., genetic underpinning) (58%) is accounted for by (or share with) those underlying the default network connectivities. Furthermore, the proportion of the phenotypic correlation between head motion and default network connectivity explained by common genes is large (48% to 61%). These findings suggest that the phenotypic correlation between head motion and default network connectivity is largely accounted by common genetic underpinnings, reflecting the “genetic pleiotropy”, which occurs when a genetic factor affects more than one trait (Solovieff, et al., 2013).

We did not conducting the global signal regression (GSR) in preprocessing the r-fMRI data, because recent studies showed that the global BOLD signal is tightly coupled with neural activity (Scholvinck, et al., 2010), indicating that the global signal may contain neurobiological meaningful information (Yang, et al., 2014). Nonetheless, to exclude the potential confounding of global signal, we conducted the trivariate genetic modeling of head motion, distant iFC of default network and global signal, and found that the genetic correlations between the distant iFC of the default network and the head motion seem unchanged, after controlling their genetic overlap with global signal (Table S9).

The specific mechanism underlying the genetic correlation between head motion and default network iFC remains to be resolved. One candidate is that the genetic correlation is accounted by the propensity of introspection (Andrews-Hanna, et al., 2014). Several studies have shown that coherent default network activity or connectivity is positively correlated with the propensity for mind wandering (Mason, et al., 2007), tendency to think about the past and future (Andrews-Hanna, et al., 2010a), self-referential (Gusnard, et al., 2001; Sheline, et al., 2009) and depressive rumination trait (Hamilton, et al., 2015). It is reasonable to assume that people with hyper-connectivity of default network tend to be more engaged in an internal thinking, and therefore move less. This explanation is consistent with the notion that the iFC in the default network might represent a neurobiological trait that predisposes some individuals to head movements (Zeng, et al., 2014), as well as the conception of the intrinsic functional connectivity as intermediate phenotype linking genes to behaviors (Khadka, et al., 2013). That is, the genetically grounded ‘intrinsic’ (covert) functional connectivity of the default network may mediate an ‘extrinsic’ (overt) head motion tendency.

It is also possible that the correlated genetic component is largely driven by heritable head motion tendency. If this is true, both positively and negatively correlated iFC should be found heritable. However, we found that the genetic overlap is only significant between head motion and iFCs negatively correlated with it, not positively correlated ones. Furthermore, the head motion driven hypothesis may predict that most of the heritability of default network iFC should be accounted by head motion tendency. However, we found that, after regressing out the effect of head motion, the heritability of residual variances of default network iFC were only minimally reduced and still significant (Table S10); in contrast, the residual variance of head motion after regressing out the distant iFC of the PCC/Precuneus became non-significant (0.15, 95% CI: 0.00-0.33). Therefore, while someone may argue that the head motion probably drives the genetic overlap observed in this study, the motion driven effect seems not to be the dominate mechanism. More studies are needed to further clarify the underlying mechanism in future.

Our findings have important implications for analyzing and interpreting functional imaging data of default network connectivity in neuropsychiatric disorders and behavioral correlates. The findings of decreased iFC within the default network in neuropsychiatric patients (e.g., autism, attention deficit hyperactivity disorder, etc.)(Andrews-Hanna, et al., 2014; Broyd, et al., 2009; Minshew and Keller, 2010; Posner, et al., 2014; Whitfield-Gabrieli and Ford, 2012) – who usually move more compared to the healthy controls (Deen and Pelphrey, 2012; Kong, et al., 2014; Wylie, et al., 2014) – may suggest that the weaker functional connectivity in patients predisposes them to move more in the scanner. In other words, both head motion tendency and default network connectivity could be genetically linked biomarkers of psychiatric disorder. Thus, simply treating head motion as an artifact or a confounding factor may subvert genuine neuronal effects (Kong, et al., 2014; Pujol, et al., 2014; Zeng, et al., 2014).

Some solutions to separate the artifactual component from the neuronal component of head motion based on fMRI data have been proposed (Power, et al., 2015), such as multiple-echo acquisitions (Kundu, et al., 2013). We propose that another strategy could include head motion estimates as variables of interest; rather than as confounding factors or nuisance variables. For example, when comparing default network functional connectivity between patients and controls, a multivariate analysis of variance with functional connectivity and head motion parameters as co-dependent variables could be employed to examine group differences. Similarly, multivariate regression analysis with the functional connectivity and head motion metrics as co-independent variables could be used to predict the severity of psychiatric symptoms or clinical status of psychiatric disorders.

Our results should be interpreted with several clarifications. First, our results may not be against the biasing effect of head motion on iFC measures and the corresponding motion correction. Although we focus on common genetic influences on default network connectivity and head motion, we also noted that the estimates of environmental contribution were also significant (Table S6), but the magnitude was smaller compared to those of genetic contribution. The motion artifact may partly underlie the environmental correlation. Here, we emphasize the genetically mediated trait component underlying the motion-iFC correlation, because it has long been ignored. Second, we used the distant degree centrality indicating the functional connectivity, therefore our results should not be extrapolated to other connectivity measures without careful examination. Third, the potential confounding of physiological processes such as respiratory and cardiac rates was not excluded in this study. However, the influence of respiration on iFC can be minimized following application of the preprocessed steps(Van Dijk, et al., 2010), as used in the study. Finally, while it is impossible to exclude all potential third confounders that may bias data quality, we confirmed the genetic correlation between head motion and default network iFC, after controlling for the genetic confounding of two data quality indicators (e.g., signal-to-noise, SNR) (see Table S11 and S12).

In conclusion, this twin study advances our understanding of the relationship between head motion and iFC of the default network. In contrast to the common view that head motion distorts iFC, our results suggest that a component of the correlation between head motion and default network iFC is mediated genetically. These findings have fundamental implications for interpreting individual differences in default network connectivity such as aberrant functional connectivity in neuropsychiatric disorders and brain-behavior associations.

## Acknowledgments

The authors thank the twins for their participating this study and thank Jie Zhang and other staffs of the BeTwiSt at the Institute of Psychology, Chinese Academy of Sciences (CAS) for their recruiting these twins. The authors gratefully acknowledge Yun Wang, Liuqing Yang, Yu Zheng and Chao-Gan Yan from Institute of Psychology, CAS for their assisting data collection and comments on early version of the manuscript. The authors acknowledge Kaibin Xu, Yue Cui from Institute of Automation, CAS for their technical assistance. The authors thank Chris Freemantle in Wellcome Trust Centre for Neuroimaging for providing extensive computational resources for the distance-dependent centrality computation. The authors acknowledge Jesper Duemose Nielsen in University of Chinese Academy of Sciences and Peter Zeidman in Wellcome Trust Centre for Neuroimaging for language editing.

### Conflict of interest

None.

